# Molecular communication enables cooperative genomic RNA replication in all-aqueous droplet colonies

**DOI:** 10.64898/2026.08.15.744999

**Authors:** Hidekazu Sono, Keiji Murayama, Kensuke Ueda, Norikazu Ichihashi, Ryo Mizuuchi

## Abstract

Multicellular organization enables biological functions to be distributed among specialized cells and coordinated through intercellular communication. Integrating this organizational principle with genome replication would link functional division of labor to the propagation of genetic information. Here, we show that genomic RNAs with complementary functions can cooperatively replicate across communicating artificial compartments. We constructed multicell-like colonies from all-aqueous droplets formed by phase separation of two incompatible polymers and stabilized at their interfaces by liposomes and amyloid-like proteins. The droplets assembled spontaneously while remaining permeable to protein-sized macromolecules. Two genomic RNAs encoding a replication enzyme and a metabolic enzyme were distributed in distinct colony-forming droplets and cooperatively replicated through cell-free translation and reciprocal molecular communication. These findings establish that genome replication can be collectively supported by communicating artificial compartments and provide a route toward multicell-like systems that coordinate spatially distributed genetic functions.

## Introduction

Bottom-up synthetic biology aims to reconstruct cellular functions using non-living materials to better understand their mechanisms and develop new biotechnologies (1–5). Artificial cells exhibiting life-like properties with a minimal set of components can also serve as model protocells for investigating the origins of life (6–8). A growing trend in the field is the creation of artificial multicellular systems to explore intercellular organization and communication, thereby displaying higher-order properties (5, 9–11). The assembly of individual artificial cells forms complex chemical reactors that enable spatiotemporal control of functions and scalable information processing (12), broadening the potential applications of artificial cells. It has also been proposed that cellular assemblages, or colonies, could represent an intermediate stage of protocell evolution, offering mechanical stability and facilitating intercellular chemical communication (13, 14). Inspired by these ideas, previous studies have demonstrated the assembly of multiple artificial cells in various forms, primarily using lipid-based vesicles (15–20) or water-in-oil droplets (21–24), but also polymersomes (25), proteinosomes (26, 27), and all-aqueous (water-in-water) droplets formed via liquid–liquid phase separation (LLPS) (28–31). Furthermore, to enable intercellular communication, some studies have incorporated gene expression machinery (21, 22, 24), which underlies most biological processes in living cells and is therefore crucial for the development of artificial cells (1–5). While these efforts have demonstrated gene-directed, regulated chemical communication and/or the expression of fluorescent proteins in artificial multicellular systems, implementing more complex biological phenomena remains challenging.

One of the fundamental cellular processes is genome replication, which underlies autonomous proliferation and Darwinian evolution across all known life forms. By employing cell-free expression systems, previous studies have demonstrated the replication of artificial genomic DNA and RNA using their self-encoded proteins within various synthetic cell forms, including cell-sized liposomes (32, 33), water-in-oil droplets (34–36), and LLPS droplets (37, 38). However, to the best of our knowledge, these systems have been limited to individual, noninteracting compartments, and genome replication has not yet been implemented in an artificial multicellular system. In such a system, molecular communication could coordinate complementary functions distributed among different artificial cells. If these functions jointly supported replication of the genomes encoding them, this would represent a step toward greater autonomy at the multicellular level. Whether such cooperative genome replication can be achieved through communication between artificial cells remains unknown.

Among the various artificial cell types, all-aqueous droplets offer unique advantages, such as the selective concentration of biomolecules and facile molecular exchange with the environment or other droplets (39, 40). Their simple biophysical mechanism of formation has also positioned all-aqueous droplets as candidates for primitive cells (41, 42). Among the different all-aqueous systems, aqueous two-phase systems (ATPSs), which typically consist of two incompatible polymers, have been particularly recognized for their wide range of biotechnological applications (43). Previous studies have shown that ATPSs support various enzymatic and catalytic reactions (44–47), including gene expression and associated genomic DNA and RNA replication (37, 38). Although all-aqueous droplets are generally unstable due to low interfacial tension and tend to coalesce, their water–water interfaces can be stabilized by various colloidal components, including polymer or protein particles (28, 48), liposomes (49, 50), and amyloid fibrils (51), while often allowing molecular exchange with the environment. Moreover, a stabilized ATPS has been used to fabricate a multicellular structure (28), although biological processes were not implemented.

Here, we developed an ATPS-based multicellular system in which molecular communication between droplets carrying complementary RNA genomes supports cooperative genome replication. In our previous work, we combined an artificial genomic RNA with a cell-free translation system in an ATPS composed of dextran (DEX) and polyethylene glycol (PEG); within DEX-rich droplets, the genomic RNA replicated using an RNA replicase (RNA-dependent RNA polymerase) translated from itself (37). Building on this system, we stabilized the droplets by incorporating PEGylated liposomes, known to accumulate at droplet surfaces (49, 50), together with amyloid-like proteins that spontaneously form within the droplets and interact with the liposomes to construct a protein-based capsule. Once stabilized, the droplets spontaneously assembled into colonies. These colonies supported translation-coupled RNA replication (TcRR) while permitting the exchange of relatively large macromolecules, such as the replicase produced within individual droplets. Using a method to detect specific RNA replication at the single-droplet level, we demonstrated genomic RNA replication mediated by multiple inter-droplet molecular communications. In particular, two genomic RNAs encoding the replicase and a metabolic enzyme, respectively, cooperatively replicated through bidirectional molecular communication between distinct droplet populations. These findings establish all-aqueous droplet colonies as a platform for constructing artificial multicellular systems that coordinate spatially distributed genetic functions to support genome replication.

## Results

### Stabilization and colony formation of ATPS droplets compatible with TcRR

We used an ATPS composed of 1.5 wt% DEX (9–11 kDa) and 15 wt% PEG (20 kDa), which formed DEX-rich phase droplets in a continuous PEG-rich phase upon vigorous mixing. This ATPS was combined with a TcRR system comprising a reconstituted *Escherichia coli* cell-free translation system (PURE system; Tables S1 and S2) (52) and a single-stranded genomic RNA (HL2-228 (53)) that encodes the catalytic subunit of Qβ replicase (Fig. 1A). Most of the translation proteins and RNA are selectively partitioned into the DEX-rich phase droplets (37). Within the droplets, the expressed catalytic subunit assembles with elongation factors Tu and Ts (EF-Tu and EF-Ts) from the translation system to form an active replicase, which subsequently replicates the genomic RNA. As previously reported, incubation of the ATPS induces TcRR reactions but also leads to droplet coalescence (37).

**Figure 1.**
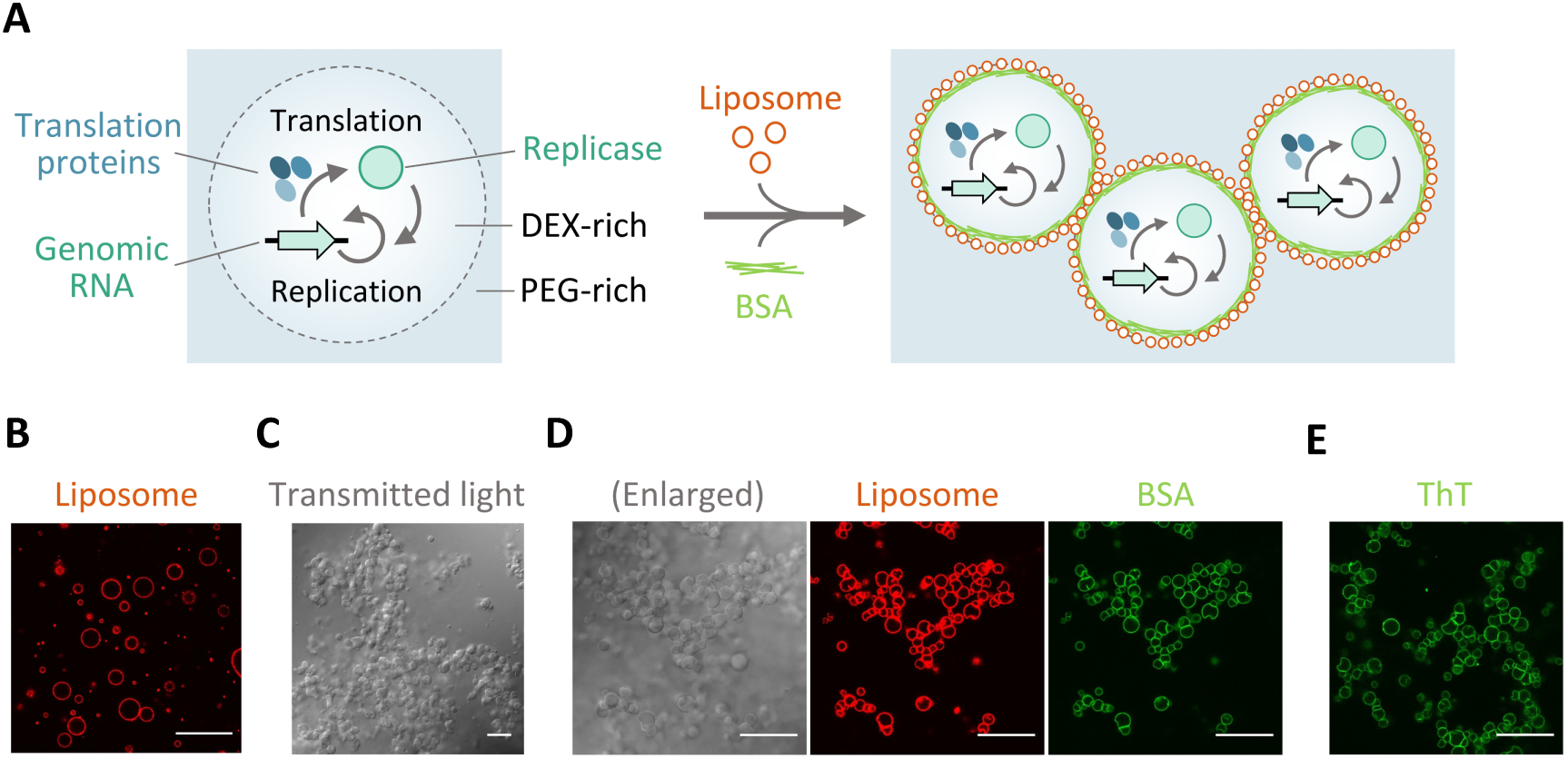
Formation of droplet colonies encapsulating the TcRR system. (**A**) Schematic of ATPS droplet colony formation mediated by liposomes and BSA. The RNA and protein components of the TcRR system spontaneously partition into the ATPS, and TcRR reaction occurs within the droplets. (**B**) Confocal images showing the accumulation of ATTO 565-labeled liposomes (0.75 mg/mL) at the ATPS droplet interface. (**C**) Transmitted light image showing droplet colony formation after incubation with BSA (1 mg/mL) at 37 °C for 4 h. (**D**) Enlarged view of droplet colonies. Left, transmitted light; middle, liposome fluorescence (ATTO 565); right, BSA fluorescence (fluorescein). (**E**) Confocal image showing ThT fluorescence after incubation at 37 °C for 4 h. Scale bars, 30 μm.

To stabilize the DEX-rich phase droplets, we introduced liposomes approximately 140 nm in diameter (Fig. S1), mainly composed of negatively charged phospholipids and PEGylated phospholipids. The liposomes primarily accumulated at the droplet interfaces (Fig. 1B), consistent with previous studies (49, 50). However, the liposome-coated droplets in our ATPS were not fully stable and underwent significant coalescence (Fig. S2) perhaps due to the relatively high ionic strength of the translation system (Table S2), a factor previously proposed to limit liposome-based droplet stabilization (49). Moreover, coating the droplets with the liposomes hindered internal TcRR reactions (Fig. 2). While the genomic RNA replicated 560-fold after 4 h incubation at 37 °C without the liposomes, the addition of the liposomes reduced replication to only 7-fold. This inhibition may have resulted from the negative charges of phospholipids, which have been shown to suppress cell-free protein translation (54). To overcome this inhibitory effect, we supplemented the ATPS with bovine serum albumin (BSA), inspired by the study that restored protein translation within negatively charged phospholipid vesicles by co-encapsulating BSA (54). We found that the addition of 1 mg/mL BSA restored the TcRR activity in the presence of the liposomes to a level comparable to that observed in their absence (Fig. 2).

**Figure 2.**
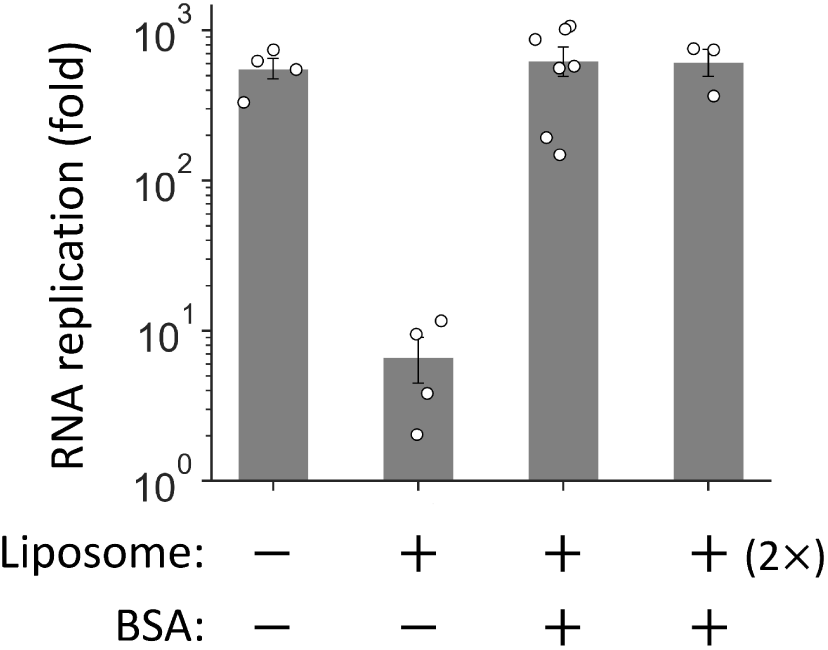
TcRR reaction in the presence of the liposomes and BSA. The TcRR system with 4 nM genomic RNA (HL2-228) was incubated at 37 °C for 4 h in the presence or absence of 0.75 mg/mL or 1.5 mg/mL (“2×”) liposomes and 1 mg/mL BSA. Replication of the genomic RNA was measured by quantitative RT-PCR. Error bars indicate mean ± SEM (*N* = 3–7, shown as individual data points).

To our surprise, we found that the addition of BSA also modulated the behavior of the liposome-coated droplets. The droplets became stable and adhered to each other without notable coalescence during 4 h incubation at 37 °C, eventually forming relatively large droplet colonies (Fig. 1A and C). To investigate how BSA stabilized the droplets, we examined the localization of fluorescein-labeled BSA in the ATPS. Before incubation (0 h), BSA molecules were evenly distributed within the droplets. However, during incubation at 37 °C, BSA gradually translocated to the droplet interface, where the liposomes accumulated, and this translocation appeared to be nearly complete after 2 h incubation (Figs. 1D, S3A, and B). This translocation was induced by incubation at 37 °C, whereas incubation at 22 °C did not alter the localization of BSA (Fig. S3C and D). Other proteins (HL2-228-encoded replicase and ribosomes), labeled with fluorescein in the same manner, remained predominantly within droplets after 2 h incubation at 37 °C (Fig. S4), suggesting that the observed translocation was likely specific to BSA.

Because BSA is known to form amyloid fibrils under diverse conditions (55–57), we hypothesized that BSA may have formed amyloids in the ATPS. To investigate this, we used Thioflavin T (ThT), a fluorescent dye whose fluorescence increases upon binding to amyloid structures. To facilitate ThT-based detection of BSA while minimizing potential interactions between ThT and RNA (58), we omitted the translation machinery, including tRNA and ribosomes, from the TcRR system. During incubation at 37 °C, ThT fluorescence intensity gradually increased around the droplet interface (Figs. 1E and S5), consistent with the localization of BSA (Fig. S3), suggesting that BSA formed amyloid-like structures that preferentially localize at the droplet interface. We also observed that BSA formed amyloid-like structures in the ATPS without the liposomes, but the droplets coalesced (Fig. S6), indicating that interactions between BSA and the liposomes contributed to droplet stabilization.

To further investigate how BSA contributed to the stabilization of liposome-coated ATPS droplets, we treated the liposomes with a surfactant after the formation of stable droplet colonies. The surfactant disrupted most of the liposomes, whereas BSA remained localized at the droplet interfaces and appeared to prevent droplet coalescence, suggesting that BSA stabilized the droplets by forming shell-like structures in the presence of liposomes (Fig. S7). Together, these results showed that liposomes and BSA converted TcRR-compatible ATPS droplets into stable, colony-forming compartments.

### Macromolecular exchange between stabilized ATPS droplets

One of the characteristic features of all-aqueous droplets is their high permeability, which allows molecular exchange between droplets. To investigate whether the colony-forming ATPS droplets remained permeable after stabilization with liposomes and BSA, we examined molecular exchange between stabilized droplets using fluorescein isothiocyanate (FITC)-labeled DEX with different molecular weights (10, 150, and 2000 kDa on average), which segregates into the DEX-rich phase (Fig. 3A). We separately prepared two populations of stabilized droplets, referred to as droplets 1 and droplets 2 for simplicity, which were coated with liposomes labeled with ATTO 565-and ATTO 647, respectively, while encapsulating FITC-labeled DEX only in droplets 1. After 2 h incubation at 37 ℃ to stabilize each population, we gently mixed them and further incubated the mixture for 4 h at 37 ℃, followed by detection of FITC signals in individual droplets. The FITC intensity in droplets 2, normalized to the mean FITC intensity in droplets 1, was used as an indicator of the efficiency of DEX exchange between droplets. Since mixing the two droplet populations slightly destabilized the system and induced leakage of the DEX-rich phase into the continuous PEG-rich phase (Fig. S8), we doubled the concentration of liposomes, which was sufficient to stabilize the droplets without inhibiting internal TcRR reactions (Fig. 2).

**Figure 3.**
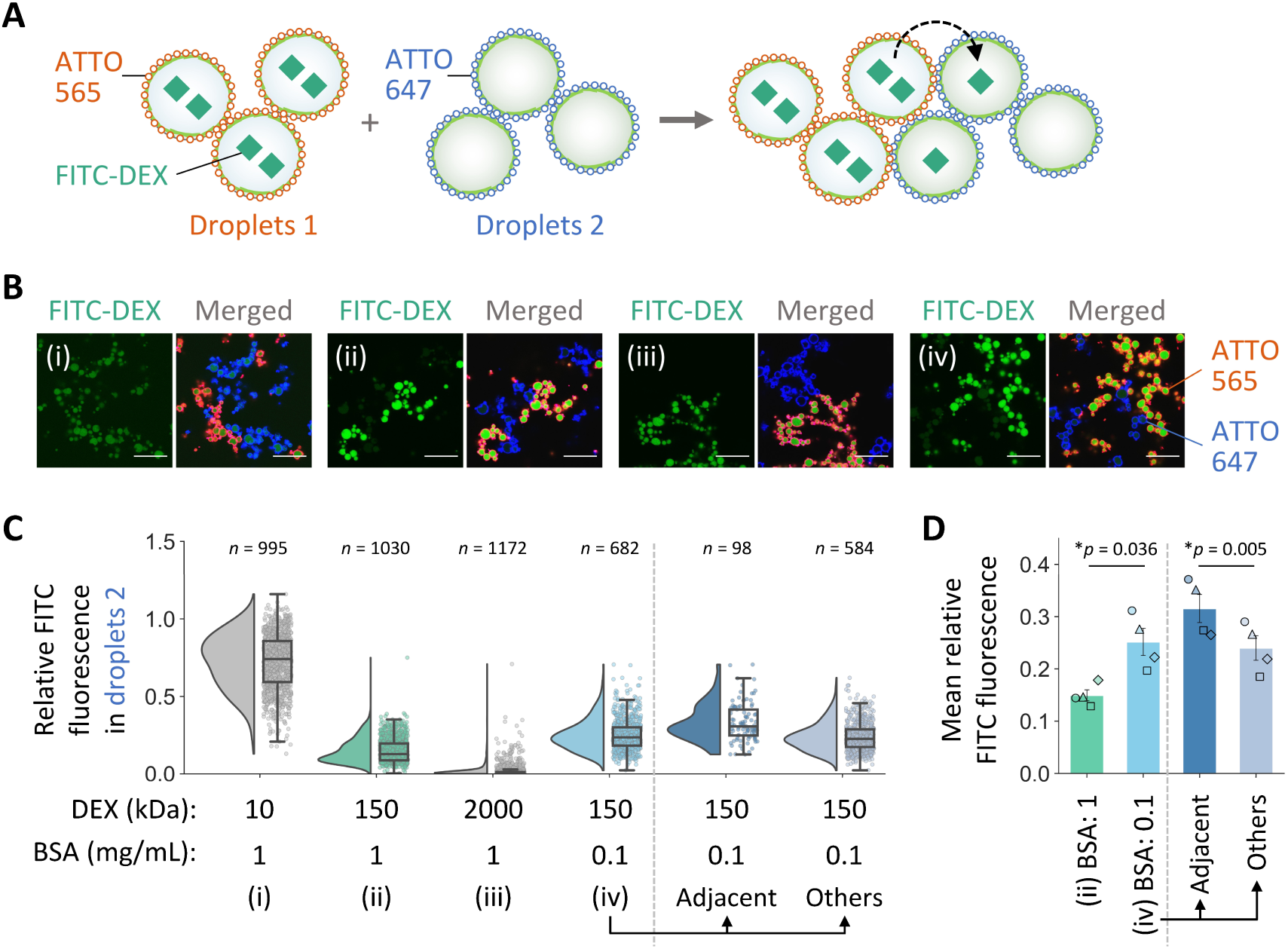
DEX exchange between stabilized, colony-forming ATPS droplets. (**A**) Schematic of the experiment. FITC-labeled DEX was encapsulated in stabilized droplets 1 and allowed to diffuse into droplets 2 during 4 h incubation at 37 °C. The two droplet populations were prepared with 1.5 mg/mL liposomes labeled with ATTO 565 or ATTO 647, which accumulated at the droplet interfaces. (**B**) Representative confocal images after 4 h incubation of the mixed droplet populations. Images (i)–(iv) correspond to the conditions shown in panel C. Left, DEX fluorescence (FITC); right, merged image of DEX fluorescence and liposome fluorescence (ATTO 565 and ATTO 647). Scale bars, 30 μm. (**C**) Distribution of FITC fluorescence intensity in droplets 2, relative to the mean intensity in droplets 1, for different combinations of DEX size and BSA concentration. “Adjacent” and “Others” represent droplets 2 in physical contact with droplets 1 in two-dimensional images and all other droplets, respectively. The number of analyzed droplets (*n*) is shown above each distribution. (**D**) Mean intensity of the distributions for 150 kDa DEX shown in panel C. Different symbols represent individual experiments (*N* = 4). Error bars show mean ± SEM. Statistical analysis was performed using a two-sided paired t-test. p-values are shown in the panel; asterisks indicate *p* < 0.05.

We observed that 10 kDa DEX efficiently diffused from droplets 1 to droplets 2, with a mean FITC intensity of 0.72 (Fig. 3B and C). Similarly, 150 kDa DEX was exchanged between the stabilized droplets, with a mean FITC intensity of 0.15, confirming that the stabilized droplets could exchange relatively large macromolecules. In contrast, the exchange of 2000 kDa DEX was undetectable in most droplets (median FITC intensity, 0.005; mean, 0.017). However, a minor fraction of droplets 2 exhibited detectable intensities, indicating that some droplets had greater permeability for macromolecular exchange.

Next, we sought to enhance the permeability of the stabilized ATPS droplets. We found that decreasing the concentration of BSA to 0.1 mg/mL, which did not affect droplet morphology (Fig. 3B), facilitated exchange of 150 kDa DEX. Under these conditions, the FITC intensities in droplets 2 became higher than those observed with 1 mg/mL BSA, with a mean intensity of 0.25 (Fig. 3C). The stabilized droplets remained largely intact for more than a month at room temperature, with only a minor fraction undergoing fusion (Fig. S9). We further hypothesized that molecular exchange could occur more efficiently between adjacent droplets than between distant droplets. To test this, the spatial localization of droplets 1 and droplets 2 was estimated using two-dimensional confocal microscopy images. We found that droplets 2 located adjacent to droplets 1 tended to exhibit higher FITC intensities (Fig. 3C and D), suggesting that nearby droplets preferentially exchange relatively large molecules.

### TcRR via inter-droplet diffusion of an encoded replicase

Given the capability of stabilized ATPS droplet colonies to exchange macromolecules, we next asked whether such molecular communication could support TcRR reactions. The observed exchange of 150 kDa FITC-DEX between droplets (Fig. 3) suggested that either the RNA replicase complex (∼138 kDa) or its catalytic subunit (∼64 kDa), translated from a genomic RNA, may diffuse from one droplet to another and initiate RNA replication. To test this, we employed a two-RNA replication system consisting of RNA 1, encoding the replicase subunit, and RNA 2, encoding nucleoside diphosphate kinase (NDK; previously designated as NDK-RNA Evo (35)) (Fig. 4A). Since TcRR using HL2-228 was severely inhibited at 0.1 mg/mL BSA (Fig. S10), we instead used a previously obtained variant with enhanced translation efficiency, R30 (59), as RNA 1. Under these conditions, RNAs 1 and 2 successfully co-replicated in the stabilized droplets, even at the reduced BSA concentration (Fig. 4B). Purified NDK was included in this experiment in addition to RNA 2, so that replication was not limited by NDK expression; bidirectional molecular communication involving NDK expression was examined later.

**Fig. 4.**
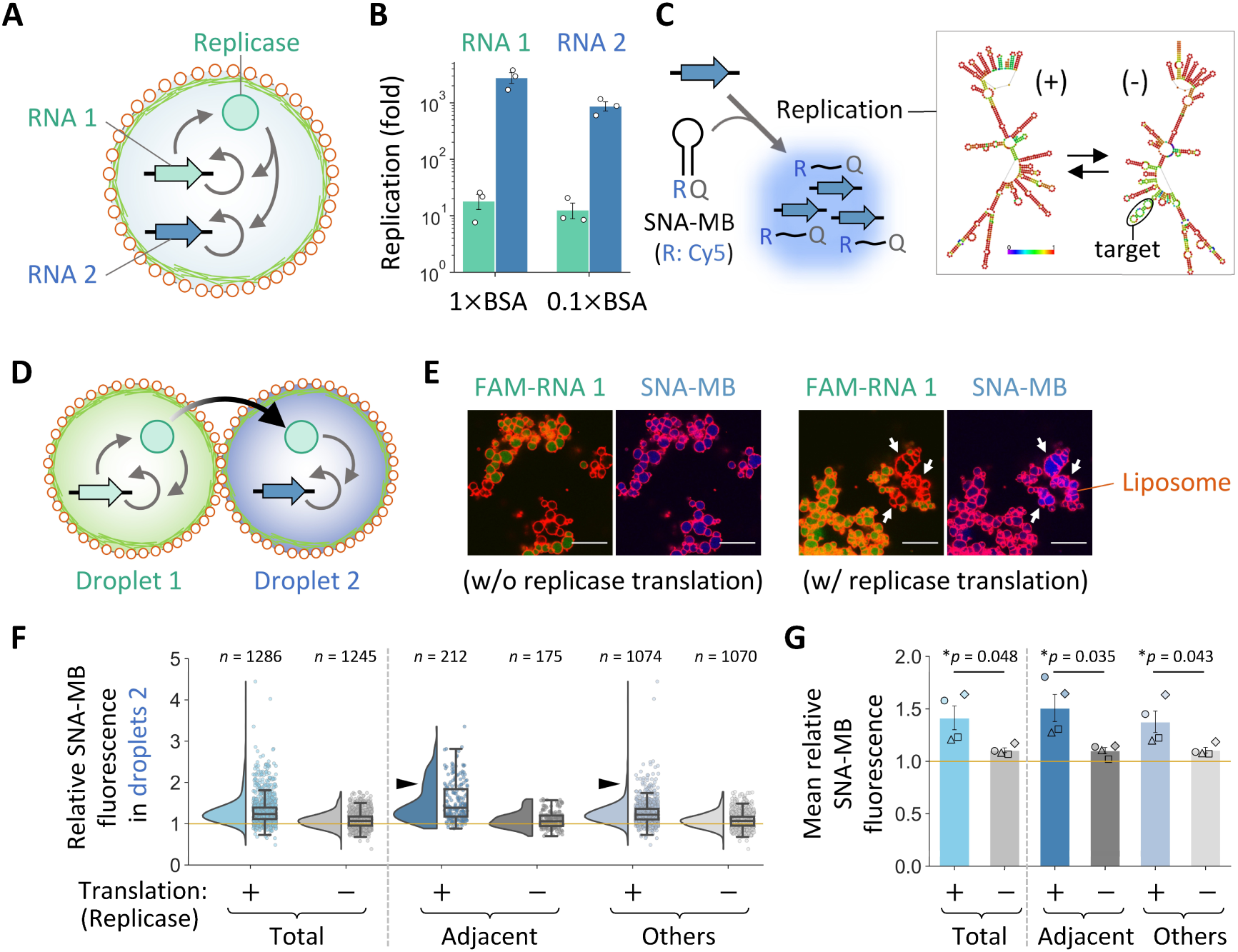
Genomic RNA replication via inter-droplet diffusion of an encoded protein in stabilized ATPS droplet colonies. (**A**) Schematic of TcRR in stabilized ATPS droplets with RNA 2. (**B**) The TcRR system with 25 nM RNA 1 and 5 nM RNA 2 was incubated at 37 °C for 4 h in the presence of 1.5 mg/mL liposomes and 1 mg/mL (“1×”) or 0.1 mg/mL (“0.1×”) BSA. Replication of the genomic RNAs was measured by quantitative RT-PCR. Error bars indicate mean ± SEM (*N* = 3, shown as individual data points). (**C**) Schematic for detection of RNA 2 replication using the SNA-MB. R and Q represent the reporter and quencher, respectively. Secondary (centroid) structures of plus-and minus-strand RNA 2 predicted by ViennaRNA (75) are shown on the right; base colors indicate base-pair probability from purple to red (more probable). (**D**) Schematic of the two droplets communicating via diffusion of the encoded replicase. The replicase translated from RNA 1 in droplet 1 can diffuse into droplet 2 to induce RNA 2 replication. The two droplet populations were prepared with 50 nM RNA 1 and 10 nM RNA 2, respectively, in the presence of 1.5 mg/mL liposomes and 0.1 mg/mL BSA. (**E**) Representative confocal images after 4 h incubation of the mixed droplet populations with or without replicase translation. Left, RNA 1 fluorescence (FAM); right, SNA-MB fluorescence (Cy5). Both images are merged with liposome fluorescence (ATTO 565). For visualization, only the MB fluorescence images were contrast-adjusted, using identical settings across images. White arrows indicate a clear increase in MB fluorescence. Scale bars, 30 μm. (**F**) Distribution of SNA-MB fluorescence intensity in droplets 2, relative to the mean intensity in droplets 1 in the NC experiment. The yellow line indicates a relative intensity of 1. The black arrowheads indicate an increased fraction of droplets 2 with relatively high fluorescence among those adjacent to droplets 1. The number of analyzed droplets (*n*) is shown above each distribution. (**G**) Mean intensity of the distributions shown in panel F. Different symbols represent individual experiments (*N* = 4). Error bars show mean ± SEM. Statistical analysis was performed using a two-sided paired t-test. p-values are shown in the panel; asterisks indicate *p* < 0.05.

To detect RNA 2 replication at the single-droplet level, we considered using a molecular beacon (MB) containing a sequence complementary to the minus (i.e., complementary) strand of RNA 2 (Fig. 4C). However, a typical MB based on single-stranded DNA (ssDNA) was not sufficiently sensitive to detect RNA 2 replication, possibly due to the stable secondary structure of RNA 2 (Fig. S11). As an alternative, we found that a similar MB based on serinol nucleic acid (SNA-MB) (60), designed to target the same region, more reliably detected RNA 2 replication (Fig. S11). This enabled us to identify droplets in which RNA 2 replication occurred.

We then prepared two droplet populations, droplets 1 and droplets 2, encapsulating RNA 1 and RNA 2, respectively, together with the TcRR system and the RNA 2-specific SNA-MB (Fig. 4D). Both populations were prepared in the presence of 1.5 mg/mL liposomes and 0.1 mg/mL BSA. To distinguish the two droplet types, fluorescein phosphoramidite (FAM)-labeled RNA 1 was co-encapsulated in droplets 1. Streptomycin, an antibiotic that inhibits protein translation, was also added to droplets 2 to ensure that any RNA 2 replication observed in these droplets could be attributed to the diffusion of the replicase rather than unintended transfer of RNA 1, if any occurred. The two droplet populations were first incubated separately for 2 h at 37 ℃ to allow droplet stabilization and the translation of the replicase from RNA 1. They were then gently mixed and further incubated for 4 h at 37 ℃. To assess RNA 2 replication, we measured the Cyanine 5 (Cy5) fluorescence intensity derived from the SNA-MB within individual droplets 2 and compared the results with negative control (NC) experiments, in which streptomycin was added to droplets 1 to preclude replicase translation. We observed an overall increase in fluorescence in droplets 2, with a significantly higher mean fluorescence intensity, indicating RNA 2 replication mediated by replicase diffusion from droplets 1 (Fig. 4E–G). Moreover, droplets 2 adjacent to droplets 1 tended to exhibit higher fluorescence intensities, as indicated by the black arrowheads in Fig. 4F, suggesting relatively efficient exchange of the replicase between nearby droplets.

We also detected elevated fluorescence intensities in a small fraction of droplets 1, although the mean fluorescence intensity was not significantly changed (Fig. S12), indicating that RNA 2 may have diffused into a subset of droplets 1. To examine potential inter-droplet RNA diffusion, we investigated the exchange of fluorescently labeled RNA 1 (2041 nt) and RNA 2 (752 nt) between droplets (Fig. S13). In most droplets, the exchange of either RNA was negligible; however, exchange of the shorter RNA 2, in particular, was evident in a small subset of droplets.

### TcCRR via bidirectional droplet–droplet molecular communication

The two-RNA system is based on translation-coupled cooperative RNA replication (TcCRR), in which RNAs 1 and 2 support each other’s replication in the absence of purified NDK (35). Using this system, we attempted to establish cooperative interactions between two droplet populations within stabilized ATPS droplet colonies (Fig. 5A). NDK, encoded by RNA 2, is a metabolic enzyme that converts cytidine diphosphate (CDP) to cytidine triphosphate (CTP), a substrate required for genomic RNA replication. By omitting both NDK and CTP while supplying CDP instead, we first examined whether RNA 2 in droplets 2 could support the replication of RNA 1 in droplets 1 through the expression of NDK. Both NDK (15.5 kDa) and the CTP produced in droplets 2 could in principle diffuse into droplets 1; however, diffusion of CTP is likely much more efficient because of its substantially smaller molecular size (37). To maximize the contribution of RNA 2 to this process, we used a highly cooperative variant, NDK-RNA N-7 (35), as RNA 2.

**Figure 5.**
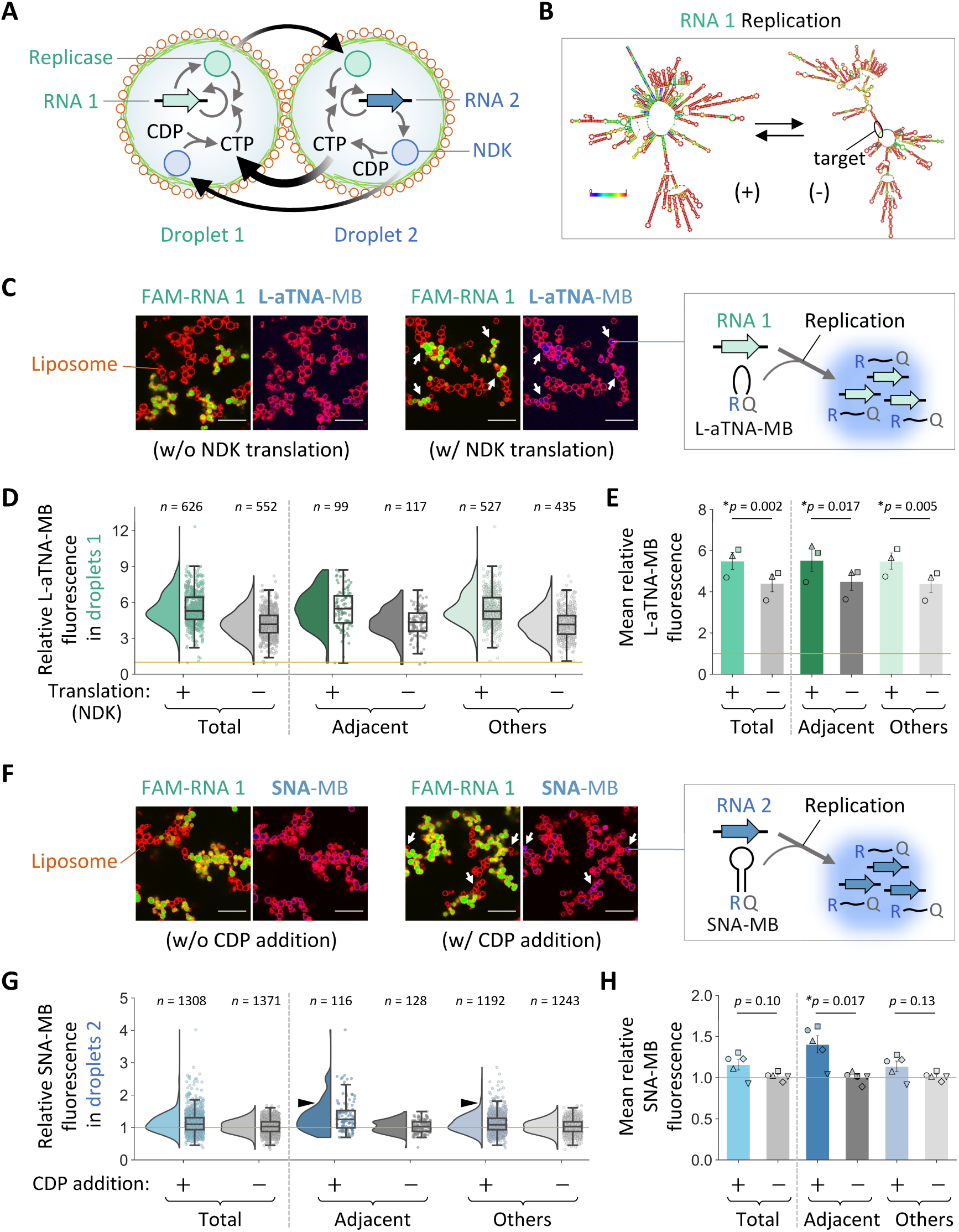
Cooperative genomic RNA replication via bidirectional droplet–droplet molecular communication in stabilized ATPS droplet colonies. (**A**) Schematic of TcCRR mediated by bidirectional molecular communication between stabilized ATPS droplets. NDK translated from RNA 2 converts CDP into CTP to support RNA 1 replication in droplets 1, while the replicase translated from RNA 1 supports RNA 2 replication in droplets 2. The two droplet populations were prepared with 100 nM RNA 1 and 10 nM RNA 2, respectively, in the presence of 1.5 mg/mL liposomes and 0.1 mg/mL BSA. (**B**) Secondary (centroid) structures of plus-and minus-strand RNA 1 predicted by ViennaRNA(75); base colors indicate base-pair probability from purple to red (more probable). The region targeted by the L-aTNA-MB is highlighted. (**C–E**) Experiment conducted using the L-aTNA-MB. (**C**) Representative confocal images after 4 h incubation of the mixed droplet populations with or without NDK translation. Left, RNA 1 fluorescence (FAM); right, L-aTNA-MB fluorescence (Cy5). Both images are merged with liposome fluorescence (ATTO 565). For visualization, only the MB fluorescence images were contrast-adjusted, using identical settings across images. White arrows indicate a clear increase in MB fluorescence. Scale bars, 30 μm. (**D**) Distribution of L-aTNA-MB fluorescence intensity in droplets 1, relative to the mean intensity in droplets 2 in the NC experiment. The yellow line indicates a relative intensity of 1. The number of analyzed droplets (*n*) is shown above each distribution. (**E**) Mean intensity of the distributions shown in panel D. Different symbols represent individual experiments (*N* = 3). Error bars show mean ± SEM. Statistical analysis was performed using a two-sided paired t-test. p-values are shown in the panel; asterisks indicate *p* < 0.05. (**F–H**) Experiment conducted using the SNA-MB, presented as in panels C–E. (**F**) Representative confocal images after 4 h incubation of the mixed droplet populations with or without the addition of CDP. (**G**) Distribution of SNA-MB fluorescence intensity in droplets 2, relative to the mean intensity in droplets 1 in the NC experiment. The black arrowheads indicate an increased fraction of droplets 2 with relatively high fluorescence among those adjacent to droplets 1. (**H**) Mean intensity of the distributions

To detect RNA 1 replication, we designed a new MB targeting the minus strand of RNA 1 (Fig. 5B) using acyclic L-threoninol nucleic acid (L-aTNA), a synthetic nucleic acid known to bind RNA tightly (61). Because strong L-aTNA–L-aTNA binding could overly stabilize the stem duplex of a conventional stem–loop beacon and slow target binding, we designed a linear beacon without a stem. We confirmed that this L-aTNA-based linear beacon (L-aTNA-MB) successfully detected RNA 1 replication at the single-droplet level (Fig. S14).

To examine cooperative interactions, we prepared droplets 1 and droplets 2 as in the experiment shown in Fig. 4D, but in the absence of NDK and CTP (Fig. 5A). The RNA 1-specific L-aTNA-MB was added to both droplet types, while FAM-labeled RNA 1 was co-encapsulated only in droplets 1 to distinguish the two droplet populations. The two droplet populations were first incubated separately for 2 h at 37 ℃ to allow droplet stabilization and the translation of the replicase and NDK from RNA 1 and RNA 2, respectively. They were then gently mixed, and CDP was simultaneously added to initiate CTP synthesis and RNA replication. Streptomycin was also added at the same time to prevent further translation of both proteins. After an additional 4 h incubation at 37 ℃, we measured the Cy5 fluorescence intensity derived from the L-aTNA-MB within individual droplets 1 to assess RNA 1 replication. These values were compared with those from negative control (NC) experiments, in which streptomycin was added to droplets 2 at the beginning to preclude NDK translation. We observed an overall increase in L-aTNA-MB-derived fluorescence in droplets 1, with a significantly higher mean fluorescence intensity, suggesting that CTP became available in droplets 1 and induced RNA 1 replication (Fig. 5C–E). As expected from the small size of CTP, droplets 1 exhibited similar fluorescence intensity distributions regardless of their spatial localization, whether adjacent to droplets 2 or not (Fig. 5D). The fluorescence increase was negligible in droplets 2, indicating that RNA 1 replication was largely limited to droplets 1 (Fig. S15A and B). We note that droplets 2 in the NC experiments also showed reduced yet measurable L-aTNA-MB-derived fluorescence compared with droplets 1 (Figs. 5D and S15A), likely because the translation protein mixture contains low levels of contaminating NDK-like activity (62).

Next, we investigated whether the replicase expressed in droplets 1 could simultaneously support RNA 2 replication through inter-droplet diffusion. To this end, we performed the same experiment by replacing the L-aTNA-MB with the RNA 2-specific SNA-MB, and assessed RNA 2 replication within individual droplets 2. Compared with NC experiments, in which CDP was not supplied, overall SNA-MB-derived fluorescence intensities increased slightly (Fig. 5F–H). Moreover, the increase in mean fluorescence became significant for droplets 2 adjacent to droplets 1, whereas it was negligible for the other droplets, suggesting that preferential exchange of the replicase between nearby droplets induced RNA 2 replication (Fig. 5G and H). As in the previous experiment (Fig. S12), a fraction of droplets 1 also showed enhanced fluorescence, suggesting that some RNA 2 diffused into droplets 1 (Fig. S15C and D). Overall, these results demonstrated cooperative genomic RNA replication mediated by bidirectional droplet–droplet molecular communication.

## Discussion

In this study, we assembled all-aqueous droplets into multicell-like colonies capable of genomic RNA replication through molecular communication. The stabilization of DEX/PEG ATPS droplets with liposomes and amyloid-like proteins led to spontaneous droplet assembly. The amyloid-like proteins accumulated on negatively charged liposomes (Fig. 1D), consistent with previous observations using giant unilamellar vesicles (63), and were likely involved in droplet–droplet adhesion (Fig. S7), reminiscent of aggregation and biofilm formation in living cells (64, 65). A unique property of the stabilized ATPS droplets is their ability to exchange not only small molecules but also protein-sized macromolecules (Fig. 3), a feature demonstrated in only a limited number of artificial cell systems (66, 67). This property expands the versatility of artificial cells and more closely mimics protein secretion-mediated communication in multicellular organisms. Moreover, this molecular communication appeared to be enhanced between adjacent droplets, while the extent of macromolecular exchange can be regulated simply by adjusting the BSA concentration (Fig. 3C and D). Leveraging these capabilities, we developed a colony composed of two droplet types, in which distinct genomic RNAs cooperate in their replication by expressing either a replication enzyme or a metabolic enzyme (Fig. 5). This type of droplet colony, in which spatially distributed genomes encode complementary functions that jointly support their replication, provides a new platform for developing multifunctional artificial cell systems.

The minimal components and functions of the TcRR and TcCRR systems incorporated into the droplets can be regarded as models of primitive genetic systems. Using these systems, previous studies have demonstrated continuous genomic RNA replication and Darwinian evolution within water-in-oil droplets, providing insights into the evolution of complex, multifunctional systems (35, 53, 62). The droplet colonies described here extend this framework from individual compartments to communicating artificial cell populations and may therefore represent a step toward demonstrating Darwinian molecular evolution at the level of artificial multicellularity. Pursuing this direction could enable investigation of the minimal requirements for higher-order evolutionary processes, such as the emergence of multicellular organization (68) and cellular differentiation (69).

We successfully detected the replication of specific single-stranded RNA (ssRNA) genomes by targeting their minus strands with MBs composed of artificial nucleic acids (Figs. S11 and S14). Detecting replicating ssRNA genomes with MBs is challenging because of rigid RNA structures, competition from complementary genomic strands for target-region binding, and interference from RNA-binding proteins such as RNA-dependent RNA polymerases. These features are likely common to both synthetic and natural ssRNA genomes (70, 71) and may have contributed to the inability of the conventional ssDNA-based MB to detect RNA 2 replication (Fig. S11). Despite these challenges, the artificial nucleic acid-based SNA-MB and L-aTNA-MB successfully visualized specific replicating RNA sequences. Given their nuclease resistance and strong RNA-binding affinity, an SNA-MB was previously used to image mRNA expression in fixed cells without washing steps (60). These properties suggest that SNA-and L-aTNA-MBs could be adapted to monitor ssRNA replication in cellular contexts, such as viral genome replication, potentially enabling direct visualization of viral infection dynamics and genome localization (72).

Together, our results establish stabilized all-aqueous droplet colonies as systems that coordinate genetic functions across spatially separated compartments. The colonies assemble spontaneously through simple mixing, providing an accessible platform for investigating molecular communication at the multicellular level mediated by cell-free gene expression. The gene expression scheme also offers programmability; additional genes and functions, including those controlling droplet-surface properties and communication with the external environment, could be incorporated in the future. Such extensions could advance the construction of artificial multicellular systems in which communication coordinates diverse functions and supports replication of the genomes encoding them.

## Materials and Methods

### Materials

Polyethylene glycol (PEG; 20 kDa), dextran (DEX; 9–11 kDa), and FITC-labeled dextran (FITC-DEX; 10, 150, and 2000 kDa), where the indicated values represent average molecular weights, were purchased from Sigma-Aldrich. Cascade Blue-labeled dextran (Cascade Blue-DEX; 10 kDa) was purchased from Thermo Fisher Scientific. PEG, DEX, FITC-DEX, and Cascade Blue-DEX were dissolved in water and stored as 40 wt%, 40 wt%, 10 wt%, and 10 wt% stock solutions, respectively. 1,2-Dioleoyl-sn-glycero-3-phospho-(1’-rac-glycerol) (sodium salt) (DOPG), 1,2-dioleoyl-sn-glycero-3-phosphoethanolamine-N-[methoxy(polyethylene glycol)-2000] (ammonium salt) (DOPE-PEG2k), and 1,2-dioleoyl-sn-glycero-3-phosphoethanolamine labeled with ATTO 565 or ATTO 647 (DOPE-ATTO 565 or DOPE-ATTO 647) were purchased from Avanti Polar Lipids. Bovine serum albumin (BSA) was purchased from Sigma-Aldrich (A6003). Replicase was obtained from our previous study as HL2-228 replicase (53). Ribosomes used in the experiment shown in Fig. S4 were obtained from PUREfrex 2.0 (GeneFrontier). Plasmids encoding HL2-228 (53), R30 (59), NDK-RNA Evo (35), and NDK-RNA N-7 (35) were obtained in our previous studies. The ssDNA-based MB was purchased from Eurofins Genomics (Tokyo, Japan), and the SNA-and L-aTNA-based MBs were purchased from Hokkaido System Science Co., Ltd. (Hokkaido, Japan). The MB sequences are listed in Table S3.

### Preparation of RNA

The cDNA of each RNA clone was PCR-amplified from the corresponding plasmid. The obtained cDNA was digested with SmaI (Takara) and subjected to *in vitro* transcription with T7 RNA polymerase (Takara) to synthesize each RNA clone. To synthesize FAM-labeled RNA, transcription was conducted in the presence of FAM-labeled UTP (Jena Bioscience). The remaining DNA was digested with DNase I (Takara). All transcribed RNAs were purified using the RNeasy Mini Kit (Qiagen).

### Preparation of the reconstituted translation system

The composition of the reconstituted translation system is shown in Tables S1 and S2. Translation proteins and ribosomes were purified as described previously (73). Briefly, each His-tagged protein was expressed in *Escherichia coli* and purified by His-tag affinity chromatography followed by size-exclusion chromatography. For TcCRR reactions, translation proteins were further purified to minimize contaminating NDK activity. Proteins other than ribosomes were subjected to two successive rounds of affinity chromatography under stringent buffer conditions, whereas ribosomes were subjected to two successive rounds of ultracentrifugation in stringent buffer, as described previously (62).

### Preparation of an ATPS containing the TcRR system

An ATPS was prepared according to a previous study (37). Briefly, DEX was first mixed with a reconstituted translation system, each RNA, and any other nucleic acids and proteins, followed by the addition of PEG. The concentrations of DEX and PEG were 1.5 wt% and 15 wt%, respectively. The DEX/PEG solution was vigorously vortexed to obtain an ATPS, in which DEX-rich phase droplets were dispersed in a continuous PEG-rich phase.

### Preparation of liposomes

To prepare liposomes, a 1 mg lipid mixture of 96.9 mol% DOPG, 3.0 mol% DOPE-PEG2k, and 0.1 mol% DOPE labeled with ATTO 565, or 647 was dissolved in chloroform and placed in a glass tube. After the lipids were dried, the resulting lipid films were hydrated with 100 μL of the PEG-rich phase of an ATPS lacking translation proteins and RNA clones to prepare a solution with a lipid concentration of 10 mg/mL. The hydrated lipid solution was sonicated for 12 s in total at ∼10 W on ice to generate liposomes. The size distribution of the liposomes was analyzed by dynamic light scattering using a Zetasizer ZSP (Malvern Panalytical) at Katayama Chemical Industries.

### Stabilization and assembly of ATPS droplets

Liposomes were added to an ATPS containing the translation system, BSA, and the indicated nucleic acids and proteins. The mixture was then vigorously vortexed. The final concentrations of BSA and liposomes were 0.1 or 1 mg/mL and 0.75 or 1.5 mg/mL, respectively, with the DEX and PEG concentrations as described above. The liposome-coated DEX-rich phase droplets were incubated at 37 °C to induce the formation of amyloid-like BSA structures, their accumulation at the liposome-coated droplet interfaces, and subsequent droplet assembly.

### Microscopic observation

Microscopic observations were performed using an FV3000 confocal laser scanning microscope (Evident) with a 40 × or a 100 × oil-immersion objective at room temperature. Cascade Blue was excited at 405 nm and imaged at 430– 470 nm; FAM, FITC, and ThT were excited at 488 nm and imaged at 500–520 nm; ATTO 565 was excited at 561 nm and imaged at 570–600 nm; and Cy5 and ATTO 647 were excited at 640 nm and imaged at 650–750 nm. To facilitate droplet segmentation during image processing, ATPS droplets for microscopic observation were prepared with 0.0000375 wt% Cascade Blue-DEX.

### Image processing

An overview of the image processing workflow is shown in Fig. S16. Stabilized ATPS droplets were segmented using Cellpose v3.1.0 with the pretrained cyto3 model (74). Liposome fluorescence images (ATTO 565 or ATTO 647) were used as the primary segmentation channel, and Cascade Blue fluorescence images were supplied as the auxiliary channel corresponding to the optional nuclear channel input of the cyto3 model. The resulting masks delineated the liposome-coated droplets.

Subsequent procedures were implemented using the Python packages SciPy and scikit-learn. For intra-droplet fluorescence quantification, only droplets with areas ≥100 pixels (6.2 μm^2^) were retained. To minimize contributions from non-target fluorescence near the droplet interface, fluorescence intensities were quantified using an inner droplet mask. For each segmented droplet, the shortest Euclidean distance from each intra-droplet pixel to the droplet exterior was calculated by applying scipy.ndimage.distance_transform_edt to its binary segmentation mask. Within each droplet, pixels with distance values above the 75th percentile were retained as the inner mask, corresponding approximately to the innermost 25% of the segmented droplet area. Using the resulting inner droplet masks, fluorescence intensities were quantified from the original, unprocessed images. Before quantification, objects whose mean liposome fluorescence intensity was ≥95% of the maximum possible pixel intensity, indicative of substantial liposome aggregation, were excluded. To reduce variation associated with differences among imaging fields or position-dependent intensity variation within images, only the top 75% of droplets with the highest Cascade Blue fluorescence intensities were retained.

In FITC-DEX and FAM-RNA exchange experiments, droplets 1 and 2 were identified based on liposome-derived ATTO 565 and ATTO 647 fluorescence, respectively. In TcRR and TcCRR experiments involving inter-droplet molecular diffusion, droplets were classified as droplets 1 or 2 based on FAM-R30 (RNA 1) fluorescence intensity. Log-transformed FAM intensities were clustered into two groups using k-means clustering implemented in sklearn.cluster.KMeans, with the higher-and lower-intensity clusters designated as droplets 1 and 2, respectively. To minimize potential misclassification near the cluster boundary, the 1% of droplets with the lowest intensities in droplets 1 and those with the highest intensities in droplets 2 were removed.

For background correction, Cascade Blue-DEX fluorescence images were thresholded using Otsu’s method to generate droplet-region masks. These masks were expanded by three iterations of binary dilation using scipy.ndimage.binary_dilation, and pixels outside the dilated masks were used as the background region. The mean fluorescence intensity in this region was calculated for each image and subtracted from the corresponding measurements of FITC-DEX, FAM-RNA, SNA-MB, and L-aTNA-MB fluorescence. To correct residual droplet-associated background in FITC-DEX and FAM-RNA exchange experiments, negative control experiments lacking the respective fluorescent molecules were performed, and the mean droplet fluorescence intensity measured in these controls was subtracted from the corresponding measurements. For each image, corrected FITC-DEX and FAM-RNA fluorescence intensities in droplets 2 were normalized to the corresponding mean fluorescence intensities in droplets 1. To quantify the SNA-MB-and L-aTNA-MB-derived fluorescence, intensities were normalized to the corresponding values in negative control experiments, as described in the text.

Adjacency between droplets from different populations was determined from the minimum Euclidean distance between the outer-boundary pixels of their segmentation masks. Droplets were classified as adjacent when this distance was <1.42 pixels (0.35 μm), encompassing direct and diagonal adjacency at the pixel level.

### ThT analysis

An ATPS was prepared with 0.33 mM ThT, a translation system lacking tRNA, translation proteins, and ribosomes, together with 0, 1, or 10 mg/mL BSA and 0.75 mg/mL liposomes. The mixture was incubated at 37 °C, and ThT fluorescence intensity was measured every minute for up to 3 h using a QuantStudio 3 Real-Time PCR System (Thermo Fisher Scientific). Microscopic observation was performed as described above.

### Protein localization analysis

Fluorescently labeled BSA, replicase, and ribosomes were prepared using the Fluorescein Labeling Kit-NH_2_ (Dojindo Molecular Technologies) according to the manufacturer’s protocol. ATPS droplets were prepared in the presence of 1 mg/mL BSA and 0.75 mg/mL liposomes, together with ∼0.03 mg/mL labeled BSA, ∼0.02 mg/mL labeled replicase, or ∼0.04 mg/mL labeled ribosomes. The mixture was incubated at 37 °C for the indicated times. Microscopic observation was performed as described above.

### DEX and RNA exchange between stabilized ATPS droplets

Two populations of ATPS droplets, droplets 1 and 2, were prepared in the presence of 0.1 or 1 mg/mL BSA and 1.5 mg/mL liposomes. The liposomes coating droplets 1 and 2 were labeled with ATTO 565 and ATTO 647, respectively. For detection of DEX exchange, 0.0038 wt% of FITC-DEX with an average molecular weight of 10 kDa, 150 kDa, or 2000 kDa was encapsulated in the droplets 1. For detection of RNA exchange, 10 nM FAM-labeled RNA 1 (R30) or RNA 2 (NDK-RNA Evo) was encapsulated in droplets 1. Droplets 1 and 2 were first incubated separately for 2 h at 37 ℃, after which equal volumes of the two droplet samples were gently mixed and further incubated for 4 h at 37 ℃. Microscopic observation was performed as described above, and FITC or FAM fluorescence intensities were determined in individual droplets.

### TcRR reaction

An ATPS containing genomic RNA and the translation system was prepared in the presence or absence of BSA and liposomes, and the mixture was incubated at 37 °C for the indicated times. To determine genomic RNA concentrations after replication, an aliquot was diluted 10,000-fold in 1 mM EDTA and subjected to RT-qPCR using the One Step TB Green PrimeScript PLUS RT-PCR Kit (Takara) and primers specific to each RNA (Table S4). A dilution series of each RNA was used to generate standard curves. When detecting RNA replication by microscopic observation, 400 nM SNA-MB or 100 nM L-aTNA-MB was added to the ATPS mixture before incubation. Microscopic observation was performed as described above.

### TcRR via inter-droplet molecular diffusion

Two populations of ATPS droplets, droplets 1 and 2, were prepared in the presence of 400 nM SNA-MB, 0.1 mg/mL BSA, and 1.5 mg/mL ATTO 565-labeled liposomes. 50 nM R30 (RNA 1) and 10 nM FAM-R30 were encapsulated in droplets 1, whereas 10 nM NDK-RNA Evo (RNA 2) and 0.06 mg/mL streptomycin were added to droplets 2. For the negative control experiment, 0.06 mg/mL streptomycin was added to droplets 1 instead of droplets 2. Droplets 1 and 2 were first incubated separately for 2 h at 37 ℃, after which equal volumes of the two droplet samples were gently mixed and further incubated for 4 h at 37 ℃. Microscopic observation was performed as described above, and SNA-MB-derived Cy5 fluorescence intensities were determined in individual droplets.

### TcCRR via inter-droplet molecular diffusion

Two populations of ATPS droplets, droplets 1 and 2, were prepared in the presence of 0.1 mg/mL BSA and 1.5 mg/mL ATTO 565-labeled liposomes. A modified reconstituted translation system was used, in which NDK, myokinase, and CTP were omitted, creatine kinase was diluted 10-fold, and the UTP concentration was doubled (Tables S1 and S2). 100 nM L-aTNA-MB was included in both droplet populations to detect RNA 1 replication, whereas 400 nM SNA-MB was included to detect RNA 2 replication. 100 nM R30 (RNA 1) and 20 nM FAM-R30 were encapsulated in droplets 1, whereas 10 nM NDK-RNA N-7 (RNA 2) was added to droplets 2. For the negative control experiment detecting RNA 1 replication, 0.06 mg/mL streptomycin was also added to the droplets 2. Droplets 1 and 2 were first incubated separately for 2 h at 37 ℃, after which equal volumes of the two droplet samples were gently mixed, and 0.03 mg/mL streptomycin, 1.25 mM magnesium acetate, and 1.25 mM CDP were added simultaneously. For the negative control experiment detecting RNA 2 replication, UDP was added instead of CDP. The droplet mixture was further incubated for 4 h at 37 ℃. Microscopic observation was performed as described above, and L-aTNA-MB-or SNA-MB-derived Cy5 fluorescence intensities were determined in individual droplets.

## Data, Materials, and Software Availability

All data supporting the findings of this study are available in the main text or the Supplementary Materials. Data files required to reproduce the analyses and custom Python code used for image processing, data preparation, and visualization are available in the associated GitHub repository: https://github.com/rmizuuchi/atps-droplet-colony-communication.

## Acknowledgments

We thank Hiroyuki Noji, Takahiro Muraoka, Takashi Kanamori, Miho Yanagisawa, and Yusuke Sato for helpful discussion. This research was supported by JSPS KAKENHI (JP21H05867 and JP23H04403 to R.M.; JP20H05970, JP20H05968, JP23H00506, and JP25K00080 to K.M.), JST CREST (JPMJCR19S4 to R.M.), and JST FOREST (JPMJFR2226 to K.M.).

## Author contributions

R.M. designed the project. H.S. and R.M. performed experiments and analyzed data. K.M. prepared the SNA-and L-aTNA-MBs. K.U. and N.I. prepared the translation system. H.S. and R.M. wrote the paper with comments from K.M., K.U., and N.I.

## Competing interests

The authors declare no competing interest.

## Notes

### Competing Interest Statement

The authors have declared no competing interest.

