## Supplementary Information for "Molecular communication enables cooperative genomic RNA replication in all-aqueous droplet colonies"

**This file includes:**
Figs. S1 to S16
Tables S1 to S4

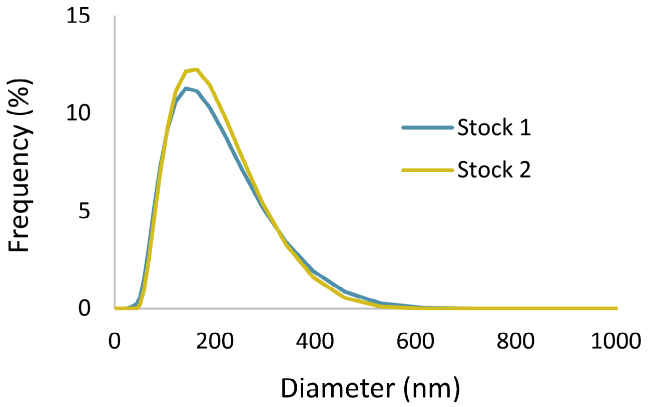


**Figure S1. Size distribution of liposomes.** The diameter of two stocks of separately prepared liposomes was analyzed by dynamic light scattering. The average diameter and polydispersity index were 139.2 nm and 0.235 for stock 1, and 139.1 nm and 0.193 for stock 2.

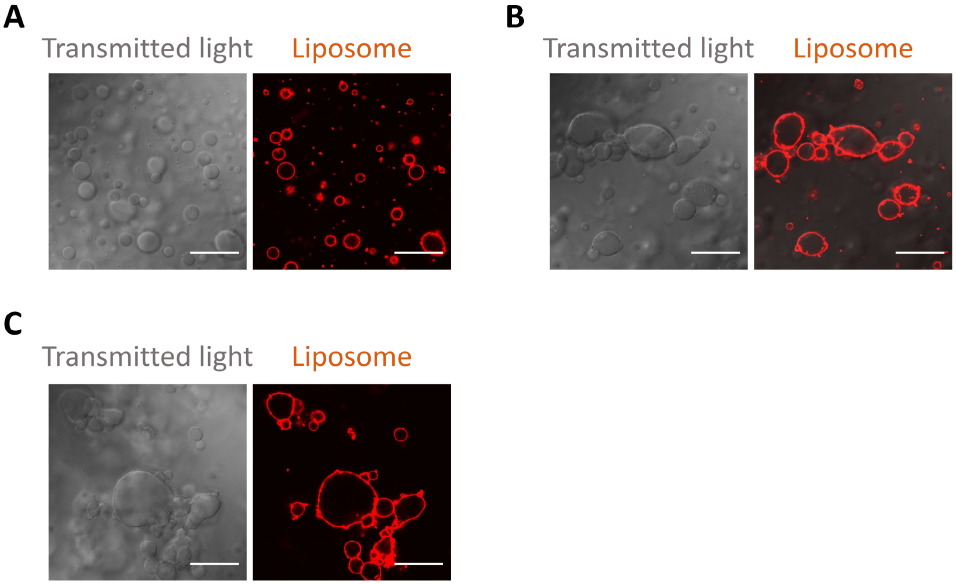


**Fig. S2. Stability of liposome-coated ATPS droplets during incubation.** Confocal images of representative liposome-coated ATPS droplets before incubation (**A**) and after incubation at 37 °C for 2 h (**B**) or 4 h (**C**). Liposomes were labeled with ATTO 565. For imaging, an aliquot of the sample was collected from the bottom phase of the solution, where droplet coalescence was most evident. Left, transmitted light; right, liposome fluorescence. Scale bars, 30 µm.

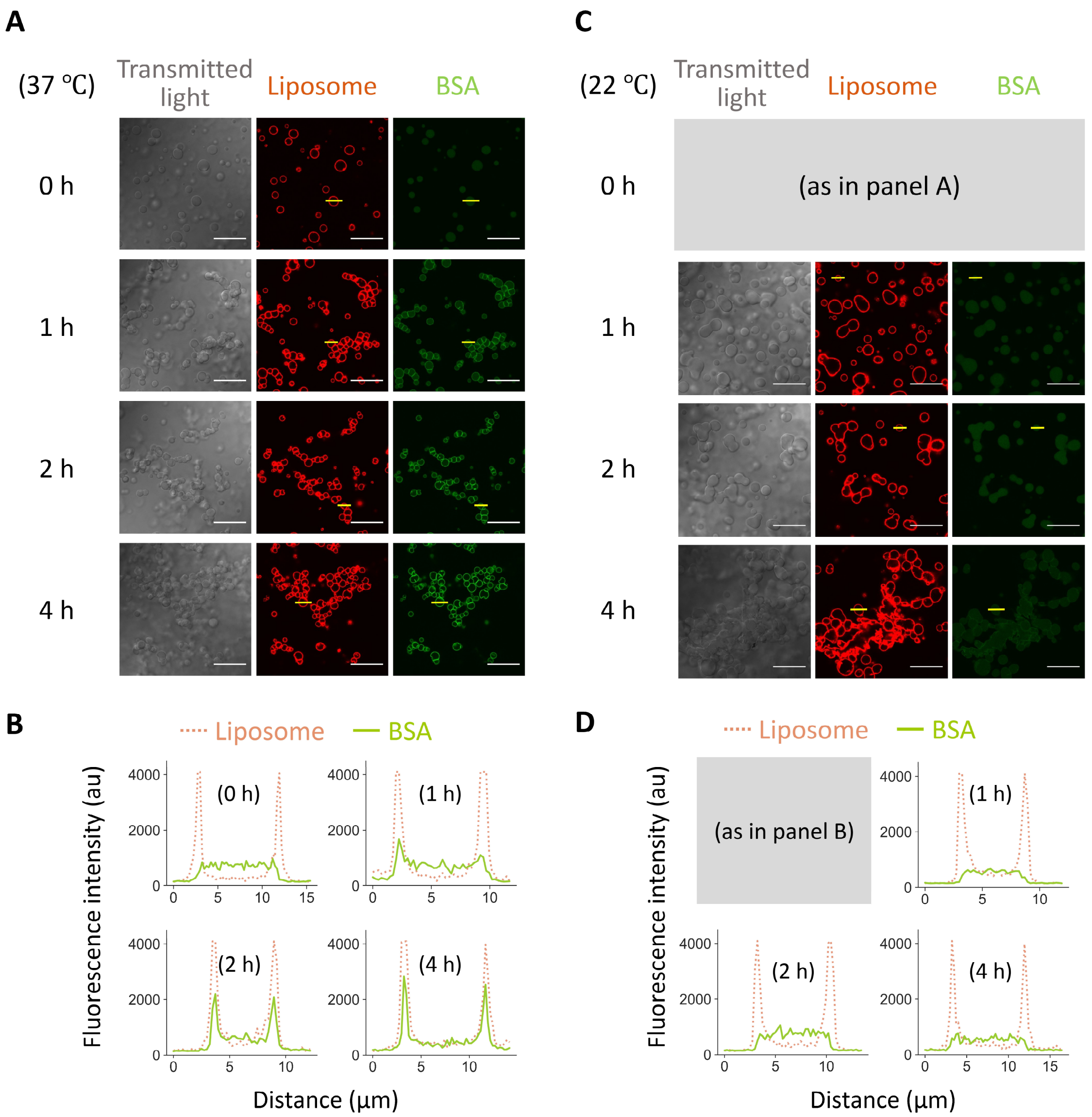


**Fig. S3. Localization of BSA in liposome-coated ATPS droplets.** (**A**) Fluorescein-labeled BSA was mixed with unlabeled BSA and incubated in the liposome-coated ATPS at 37 °C, with a total BSA concentration of 1 mg/mL. Representative confocal images after 0, 1, 2, and 4 h of incubation are shown. The images at 4 h are the same as those shown in Fig. 1D. Liposomes were labeled with ATTO 565. Left, transmitted light; middle, liposome fluorescence; right, BSA fluorescence. Scale bars, 30 μm. (**B**) Fluorescence intensity profiles along the solid yellow lines in the images in panel A. (**C** and **D**) The same analyses performed after incubation at 22 °C.

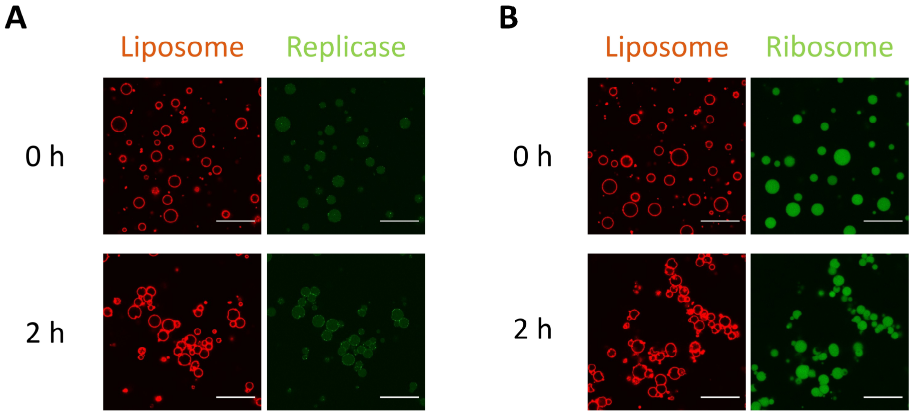


**Fig. S4. Localization of purified replicase and ribosomes in liposome-coated ATPS droplets.** Fluorescein-labeled HL2-228-derived replicase (**A**) and ribosomes (**B**) were incubated in the liposome-coated ATPS containing 1 mg/mL BSA at 37 °C. Representative confocal images at 0 and 2 h are shown. Liposomes were labeled with ATTO 565. Left, liposome fluorescence; right, protein fluorescence. Scale bars, 30 µm.

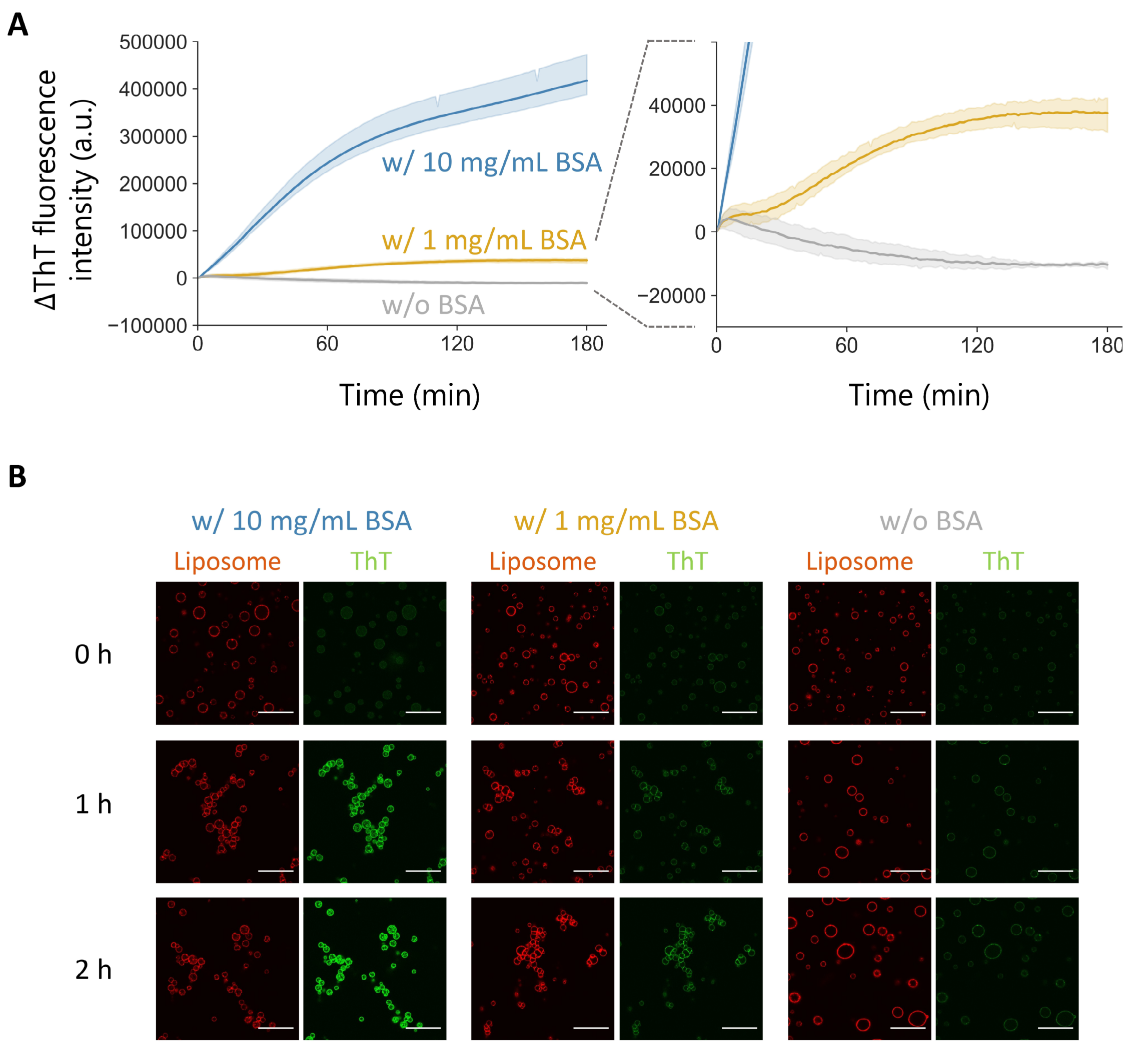


**Fig. S5. Formation of amyloid fibrils detected using ThT.** (**A**) ThT fluorescence intensity was measured during incubation of the ATPS containing 0, 1, or 10 mg/mL BSA at 37 °C. tRNA and translation proteins, including ribosomes, were omitted from the ATPS to facilitate detection. Shaded regions represent 95% confidence intervals. An enlarged view of the plot is shown on the right. (**B**) Representative confocal images of the ATPS during incubation. Liposomes were labeled with ATTO 565. Left, liposome fluorescence; right, ThT fluorescence. Scale bars, 30 μm.

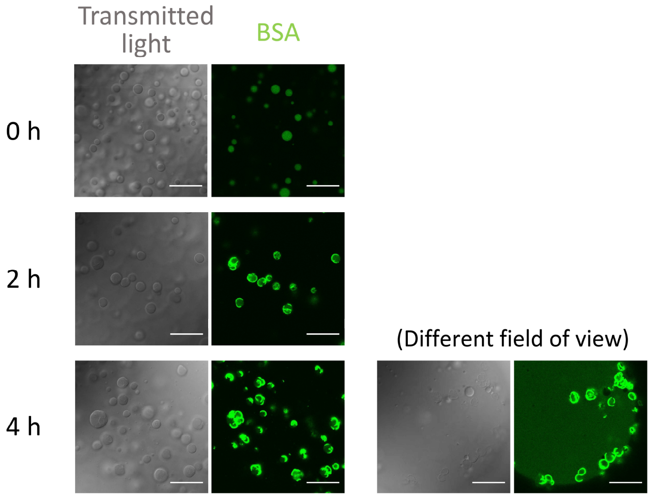


**Fig. S6. BSA fibrillization in the ATPS droplets in the absence of liposomes.** Fluorescein-labeled BSA, mixed with unlabeled BSA, was incubated in the ATPS without liposomes at 37 °C (total BSA concentration, 1 mg/mL). Representative confocal images at 0, 2, and 4 h are shown. For imaging, an aliquot of the sample was collected from the bottom phase of the solution, where droplet coalescence was most evident. Multiple fields of view at 4 h are shown. Left, transmitted light; right, BSA fluorescence. Scale bars, 30 μm.

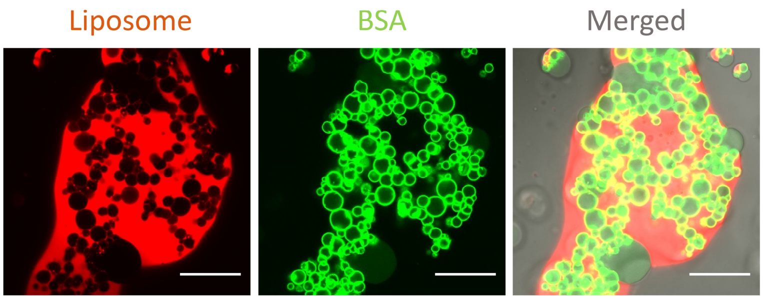


**Fig. S7. Addition of a surfactant to stabilized ATPS droplets.** ATPS droplets stabilized after 2 h incubation at 37 °C were mixed with 3% volume of Triton X-100, followed by incubation for 1 h at 37 °C. After incubation, droplets were imaged by confocal microscopy. Left, liposome fluorescence (ATTO 565); middle, BSA fluorescence (fluorescein); right, merged image of both fluorescence channels and transmitted light. Scale bars, 30 μm.

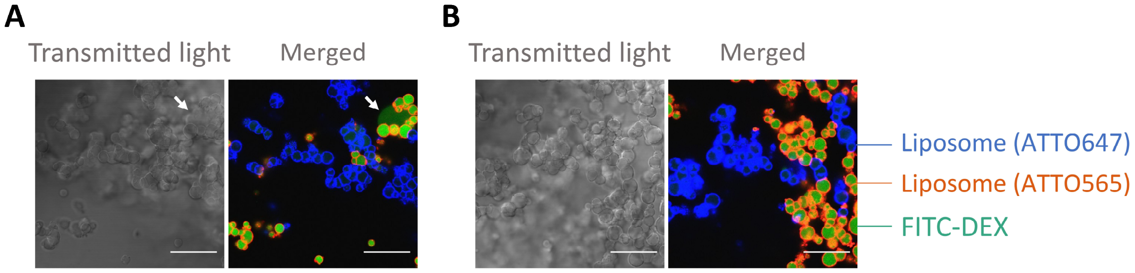


**Fig. S8. Effect of mixing two droplet populations on droplet stability.** The experiment was performed as in Fig. 3, but with 0.75 mg/mL liposomes (**A**) or 1.5 mg/mL liposomes (**B**). The average molecular weight of FITC-DEX was 150 kDa. Left, transmitted light; right, merged image of DEX fluorescence (FITC) and liposome fluorescence (ATTO 565 and ATTO 647). White arrows in panel A indicate leaked DEX-rich phases. Scale bars, 30 μm.

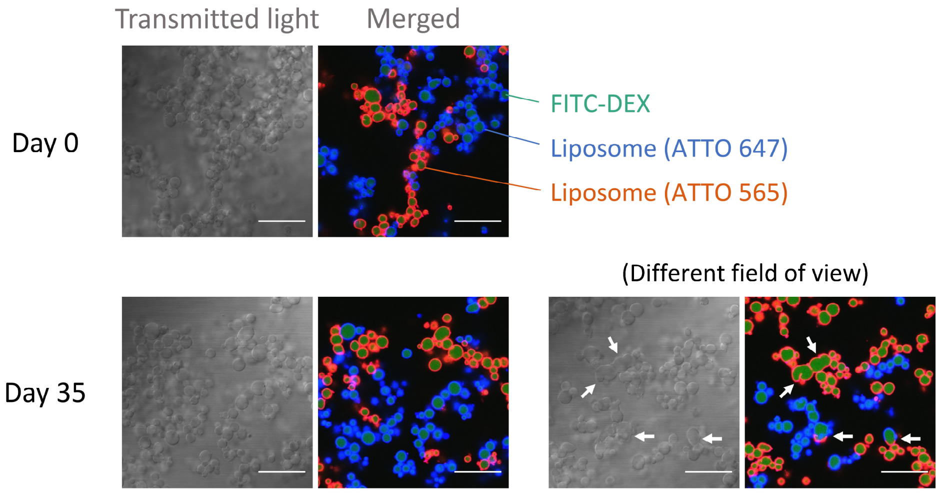


**Fig. S9. Long-term incubation of stabilized ATPS droplets.** Two droplet populations were prepared in the presence of 0.1 mg/mL BSA and 1.5 mg/mL liposomes labeled with ATTO 565 or ATTO 647, incubated separately for 2 h at 37 °C, then mixed and further incubated for 4 h (Day 0). FITC-labeled DEX with an average molecular weight of 10 kDa was encapsulated in both droplet populations. The stabilized droplets were subsequently stored at room temperature for 35 days and visualized by confocal microscopy. Confocal images obtained before (top) and after (bottom) storage are shown; multiple fields of view after 35 days are presented. Left, transmitted light; right, merged image of liposome fluorescence (ATTO 565 and ATTO 647) and DEX fluorescence (FITC). White arrows indicate droplets that apparently underwent droplet–droplet fusion. Scale bars, 30 μm.

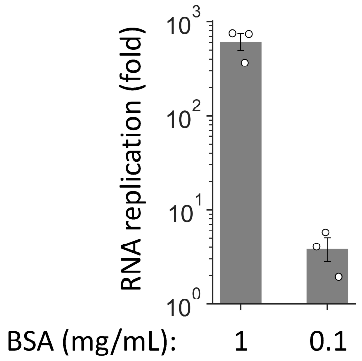


**Fig. S10. TcRR reaction in the presence of liposomes and reduced BSA.** The TcRR system with 4 nM HL2-228 was incubated at 37 °C for 4 h in the presence of 1.5 mg/mL liposomes and either 1 or 0.1 mg/mL BSA. Replication of the genomic RNA was measured by quantitative RT-PCR. The data for 1 mg/mL BSA are the same as those shown in Fig. 2. Error bars indicate mean ± SEM ($N = 3$, shown as individual data points).

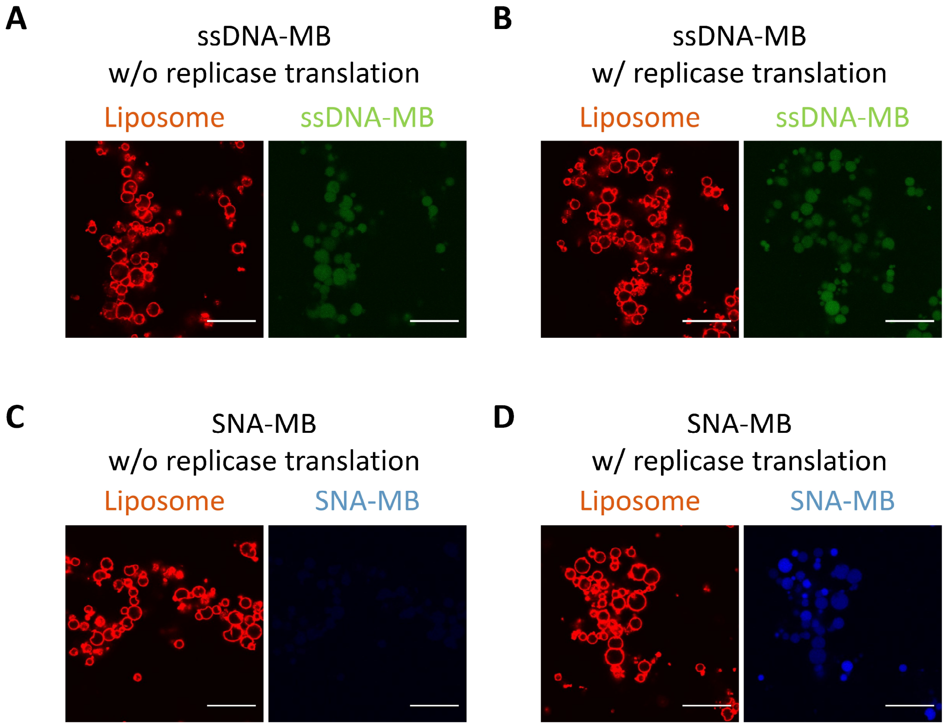


**Fig. S11. Detection of RNA 2 replication using the SNA-MB.** The TcRR system with 50 nM RNA 1 (R30) and 10 nM RNA 2 (NDK-RNA Evo) was incubated at 37 °C for 4 h in the presence of 0.75 mg/mL liposomes, 1 mg/mL BSA, and 400 nM ssDNA-MB (**A**, **B**) or SNA-MB (**C**, **D**), both targeting the same sequence region, with (A, C) or without (B, D) streptomycin to prevent replicase translation. After incubation, the droplets were visualized by confocal microscopy. Left, liposome fluorescence (ATTO 565); right, MB fluorescence (FAM or Cy5). Scale bars, 30 µm.

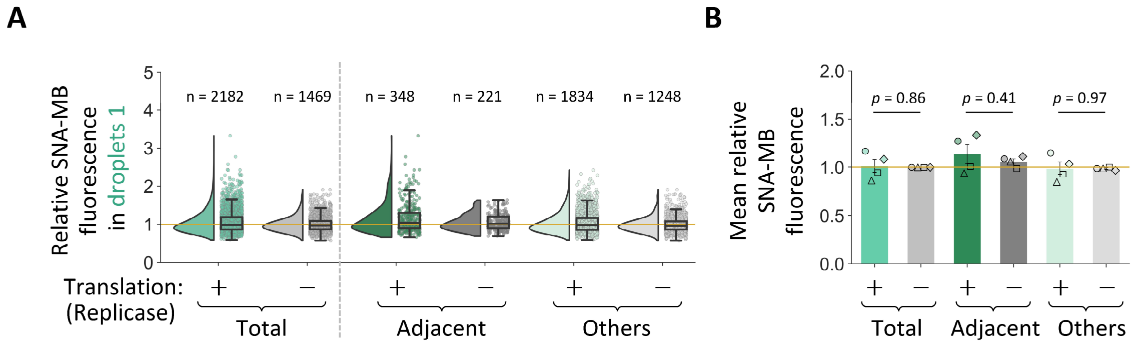


**Fig. S12. Genomic RNA replication via inter-droplet communication of an encoded protein: analysis of droplets 1.** (**A**) Distribution of SNA-MB fluorescence intensity in droplets 1, relative to the mean intensity in droplets 1 in the NC experiment. The yellow line indicates a relative intensity of 1. The number of analyzed droplets (*n*) is shown above each distribution. (**B**) Mean intensity of the distributions shown in panel A. Different symbols represent individual experiments ($N = 4$). Error bars show mean ± SEM. Statistical analysis was performed using a two-sided paired t-test. p-values are shown in the panel; asterisks indicate $p < 0.05$.

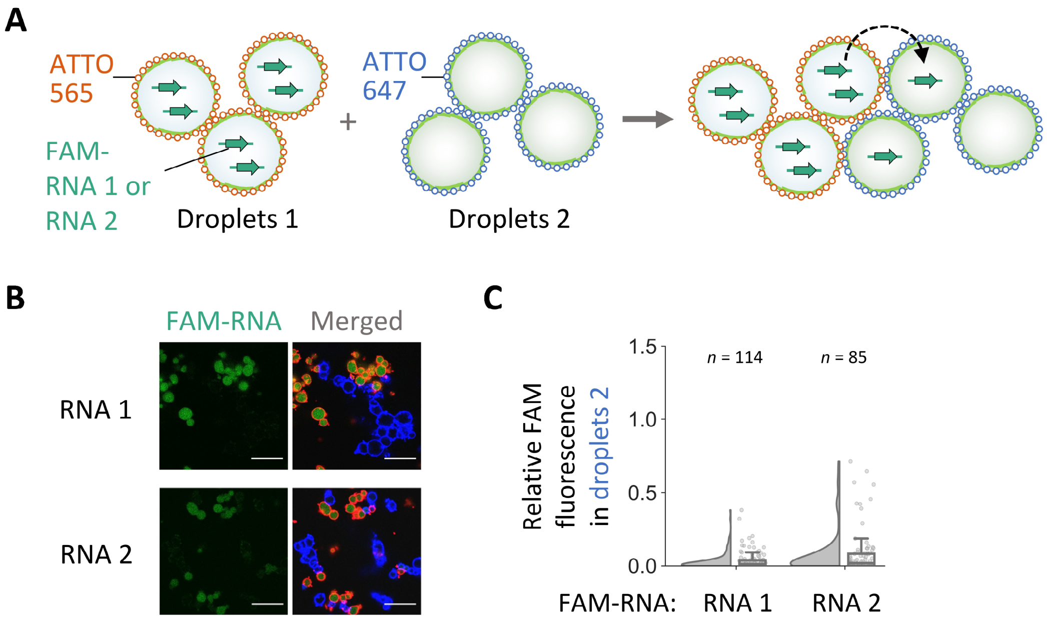


**Fig. S13. FAM-RNA exchange between stabilized ATPS droplets.** (**A**) Schematic of the experiment. FAM-labeled RNA 1 or RNA 2 (10 nM) was encapsulated in stabilized droplets 1 and allowed to diffuse into droplets 2 during 4 h incubation at 37 °C. Liposomes and BSA were used at 1.5 mg/mL and 0.1 mg/mL, respectively. Liposomes accumulating at the interfaces of the two droplet populations were labeled with different fluorophores, ATTO 565 and ATTO 647. (**B**) Representative confocal images after 4 h incubation of the mixed droplet populations. Left, RNA fluorescence (FAM); right, merged image of RNA fluorescence and liposome fluorescence (ATTO 565 and ATTO 647). Scale bars, 30 µm. (**C**) Distribution of FAM fluorescence intensity in droplets 2, relative to the average intensity in droplets 1. The number of analyzed droplets (*n*) is shown above each distribution. For RNA 1, the median and mean relative FAM intensities were 0.013 and 0.036, respectively. For RNA 2, they were 0.019 and 0.083, respectively.

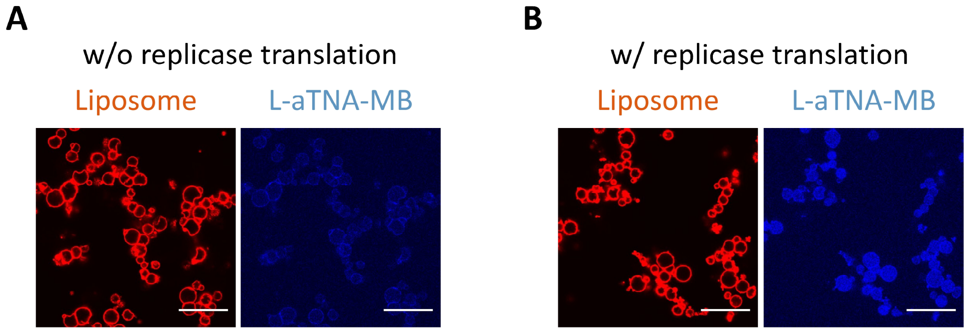


**Fig. S14. Detection of RNA 1 replication using the L-aTNA-MB.** The TcRR system with 100 nM RNA 1 (R30) was incubated at 37 °C for 4 h in the presence of 0.75 mg/mL liposomes, 1 mg/mL BSA, and 100 nM L-aTNA-MB, with (**A**) or without (**B**) streptomycin to prevent replicase translation. After incubation, the droplets were visualized by confocal microscopy. Left, liposome fluorescence (ATTO 565); right, L-aTNA-MB fluorescence (Cy5). For visualization, only the MB fluorescence images were contrast-adjusted, using identical settings across images. Scale bars, 30 µm.

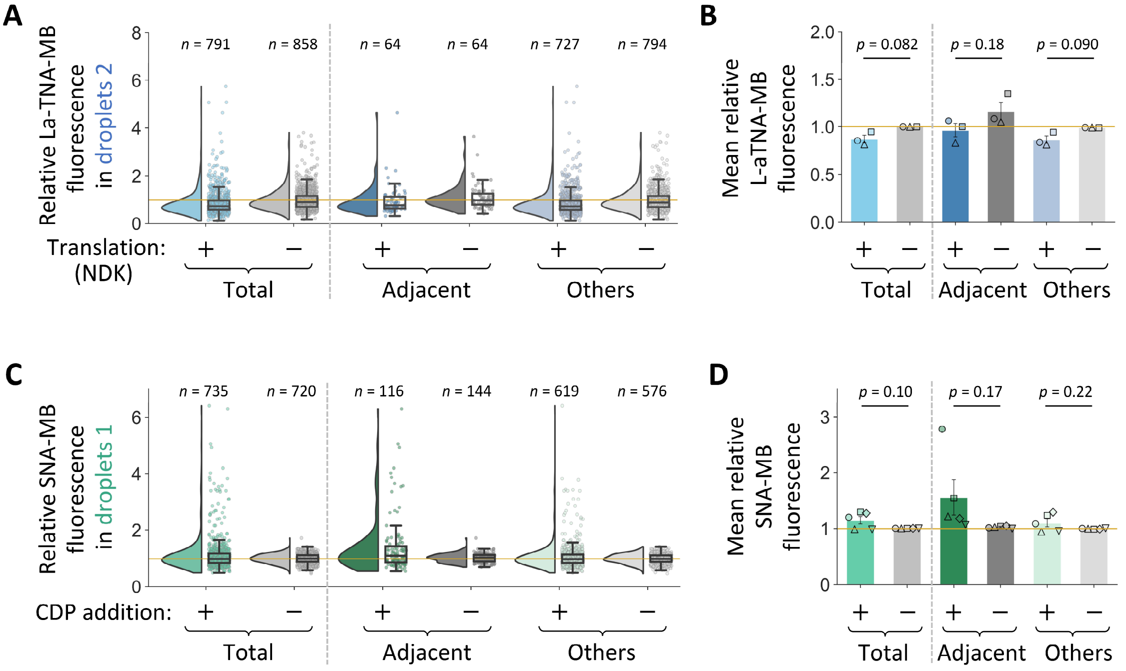


**Fig. S15. Cooperative genomic RNA replication via bidirectional droplet–droplet molecular communication: analysis of the other droplet populations.** (**A, B**) Experiment conducted using the L-aTNA-MB. (**A**) Distribution of L-aTNA-MB fluorescence intensity in droplets 2, relative to the mean intensity in droplets 2 in the NC experiment. The yellow line indicates a relative intensity of 1. The number of analyzed droplets ($n$) is shown above each distribution. (**B**) Mean intensity of the distributions shown in panel A. Different symbols represent individual experiments ($N = 3$). Error bars show mean ± SEM. Statistical analysis was performed using a two-sided paired t-test. p-values are shown in the panel; asterisks indicate $p < 0.05$. (**C, D**) Experiment conducted using the SNA-MB, presented as in panels A and B. (**C**) Distribution of SNA-MB fluorescence intensity in droplets 1, relative to the mean intensity in droplets 1 in the NC experiment. (**D**) Mean intensity of the distributions shown in panel C ($N = 5$).

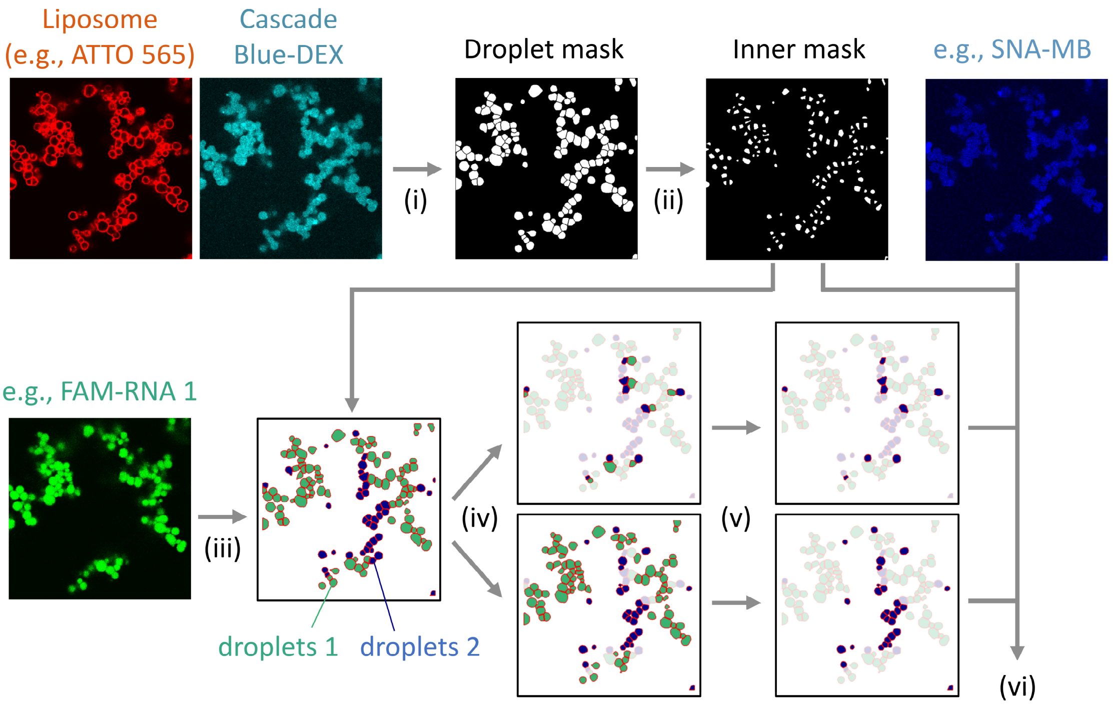


**Fig. S16. Overview of the image processing workflow.** (i) Stabilized ATPS droplets were segmented using liposome fluorescence and Cascade Blue-DEX images to generate droplet masks. (ii) Inner droplet masks were generated by thresholding the distance transform. (iii) Droplets 1 and 2 were identified based on liposome fluorescence (ATTO 565 and ATTO 647) or FAM-RNA 1 fluorescence, depending on the experiment. (iv) Adjacency between droplets 1 and 2 was determined, and (v) the relevant droplet population was selected. (vi) Fluorescence intensities of target molecules, such as SNA-MB, were quantified after background correction.

**Table S1. Protein components of the reconstituted translation system**

| Name | Concentration (nM) | Names | Concentration (nM) |
|---|---|---|---|
| Initiation factor 1 | 10000 | Isoleucyl-tRNA synthetase | 148 |
| Initiation factor 2 | 400 | Leucyl-tRNA synthetase | 16.4 |
| Initiation factor 3 | 1960 | Lysyl-tRNA synthetase | 48 |
| Elongation factor G | 440 | Methionyl-tRNA synthetase | 44 |
| Elongation factor Tu | 32000 | Phenylalanyl-tRNA synthetase | 52 |
| Elongation factor Ts | 1320 | Prolyl-tRNA synthetase | 68 |
| Release factor 1 | 19.6 | Seryl-tRNA synthetase | 31.2 |
| Release factor 2 | 19.2 | Threonyl-tRNA synthetase | 33.6 |
| Release factor 3 | 68 | Tryptophanyl-tRNA synthetase | 11.2 |
| Ribosome recycling factor | 1560 | Tyrosyl-tRNA synthetase | 60 |
| Alanyl-tRNA synthetase | 292 | Valyl-tRNA synthetase | 6.8 |
| Arginyl-tRNA synthetase | 12.4 | Methionyl-tRNA formyltransferase | 236 |
| Asparaginyl-tRNA synthetase | 168 | Myokinase* | 560 |
| Asparagyl-tRNA synthetase | 48 | Creatine kinase* | 100 |
| Cysteinyl-tRNA synthetase | 9.6 | Nucleoside diphosphate kinase* | 6.4 |
| Glutaminyl-tRNA synthetase | 24 | Pyrophosphatase | 16.4 |
| Glutamyl-tRNA synthetase | 92 | Trigger factor | 400 |
| Glycyl-tRNA synthetase | 34.4 | ATP-dependent RNA helicase HrpA | 40 |
| Histidyl-tRNA synthetase | 34 | Ribosome | 400 |

*For experiments shown in Fig. 5, myokinase and nucleoside diphosphate kinase were omitted, and creatine kinase was 10-fold diluted.

**Table S2. Other components of the reconstituted translation system**

| Name | Concentration |
|---|---|
| Tyrosine | 0.3 mM |
| Cysteine | 0.3 mM |
| 18 other amino acids | 0.36 mM each |
| tRNA mix (Roche) | 1.56 μg/μL |
| ATP | 3.75 mM |
| GTP | 2.5 mM |
| CTP* | 1.25 mM |
| UTP | 1.25 mM |
| N-2-Hydroxyethylpiperazine-N'-2-ethanesulfonic acid (pH 7.6) | 100 mM |
| Glutamic acid potassium salt | 70 mM |
| Spermidine | 0.375 mM |
| Magnesium acetate | 9.5 mM |
| Creatine phosphate | 25 mM |
| Dithiothreitol | 6 mM |
| 10-Formyl-5,6,7,8-tetrahydrofolic acid | 10 ng/μl |

*For experiments shown in Fig. 5, CTP was omitted.

**Table S3. Molecular beacons used in this study**

| Name | Target RNA | Sequence |
|---|---|---|
| ssDNA-MB | (-) RNA 2 | 5′-FAM-GCGACTGTGCTGGAAGGTGAAAATCGC-BHQ-1-3′ |
| SNA-MB | (-) RNA 2 | (*S*) Cy5-CTGTGCTGGAAGGTGAAAAACAG-BHQ-2 (*R*) |
| L-aTNA-MB | (-) RNA 1 | 3′-Cy5-TGTGAGATACTGGA-BHQ-1-1′ |

The underlined sequence regions are expected to form stem structures. The regions highlighted in red are complementary to the target RNA sequences. BHQ, Black Hole Quencher.

**Table S4. Primers used for quantitative RT-PCR**

| Name | Target RNA | Sequence |
|---|---|---|
| Primer 1 | HL2-228, R30 (RNA 1) | 5′-CAAGTATCGTAAGTTGCTGCC-3′ |
| Primer 2 | HL2-228, R30 (RNA 1) | 5′-CCGTAATCACCGGTACGTAC-3′ |
| Primer 3 | NDK-RNA Evo (RNA 2) | 5′-CGATCGTGGTTTCTGTGCTGG-3′ |
| Primer 4 | NDK-RNA Evo (RNA 2) | 5′-CCAGCGCATTTGCCGGATTG-3′ |
